# Standard methods for marking caudate amphibians do not impair animal welfare over the short term: an experimental approach

**DOI:** 10.1101/2023.09.28.560063

**Authors:** DR Daversa, E Baxter, GM Rosa, C Sergeant, TWJ Garner

**Affiliations:** La Kretz Center for California ConservaLon Science, InsLtute of the Environment and Sustainability, University of California, Los Angeles, USA; InsLtute of Zoology, Zoological Society of London, Regents Park, London, United Kingdom; Centre for Ecology, EvoluLon and Environmental Changes (cE3c) & Global Change and Sustainability InsLtute (CHANGE), Faculdade de Ciências da Universidade de Lisboa, Lisboa, Portugal; Department of GeneLcs, EvoluLon and Environment, UCL, London, United Kingdom; Unit for Environmental Sciences and Management, North-West University, Potchefstroom, South Africa

**Keywords:** Welfare Science, Marking methods, Animal Behaviour, Wildlife health, Experimental Biology, Amphibians

## Abstract

Major advancements in ecology and biodiversity conservation have been made thanks to methods for marking and individually tracking animals. Marking animals is both widely used and controversial due to the potential consequences to animal welfare, which are often incompletely evaluated before implementation. Two outstanding knowledge gaps concerning the welfare consequences of individual marking concerns their short-term behavioural impacts and the relative impacts from marking versus the handling of animals while carrying out procedures. We addressed these knowledge gaps through an experimental study of alpine newts (*Ichthyosaura alpestris*) in which we varied handling and marking procedures. Examining individual responses to handling, toe-clipping and visible implant elastomer (VIE) injection over 21 days showed that handling and marking elicited increased newt activity and hesitancy to feed compared to animals that did not get handled or marked. These effects were apparent even when animals were handled only (not marked), and marking did not further increase the magnitude of responses. Increases in newt activity and feeding hesitancy were transient; they were not observed in the weeks following handling and marking. Whereas previous studies emphasize the welfare impacts of marking procedures themselves, these findings highlight that handling alone can elicit behavioural changes with possible costs to welfare. Yet, the transient nature of behavioural responses observed here suggests that immediate costs of handling may be subsequently compensated for.

## Introduction

Confronting the global biodiversity crisis requires a critical understanding of how threatening processes impact wildlife populations. Demographic studies are the most common approach for investigating impacts, studies which frequently require the capacity to discriminate amongst individuals (Major et al. 2020). For many species, this requires handling and the application of an artificial mark. A fundamental principle of wildlife marking is the minimizing of pain and distress; the mark ideally should not significantly impair the welfare of the marked individual (e.g. Locatelli et al. 2019). Despite the widespread acceptance of prioritizing welfare (Dawkins, 2006; Hecht, 2021; Soryl et al. 2021), the impacts of capture and marking are not always tested, or at least not revisited regularly to reassess the lasting consequence of marking methods under different circumstances (Soulsbury et al. 2020). As a result, some marking techniques have gained widespread application across taxonomic groups without explicit tests of impacts on each species, or reapplication in new populations, with examples including colour-coded bird banding, fish adipose fin clipping and amphibian and reptile toe-clipping (Uglem et al 2019; Tinbergen et al 2014; Perry et al 2011).

Amphibians are the most threatened vertebrate taxon (Stuart et al., 2004; Catenazzi, 2015; Scheele et al. 2019), a conclusion largely justified through the outputs of numerous studies of population dynamics utilizing individual marking strategies (e.g. Bucciarelli et al. 2020, Storfer, 2003). Although less invasive methods are available, their use is frequently hampered by a range of limitations. For instance, individual skin patterns, while an option, often lack distinctiveness or prove suitable solely for short-term studies due to their tendency to change over time (Arntzen et al. 2004; Ferner, 2007); conversely, radio tracking, while a powerful tool, presents size restrictions, feasible only for relatively larger specimens, while bearing substantial economic costs (Ferner, 2007; Andreone et al. 2013). As a result, researchers have traditionally resorted to established methods like toe-clipping (the complete or partial removal of digits; eg. Perry et al. 2011) and, more recently, visible implant elastomer (VIE) injection (subcutaneous injection of silicone-based polymer that hardens after injection), which are attractive due to the inexpensive costs and relatively fast execution that permits large sample sizes.

Toe clipping and VIE injection, while instrumental in assessing how threatening processes impact demography, have sparked controversies concerning welfare implications (Perry et al 2011). Critical studies of their impacts have revealed real and potential impacts on individual survival. Marking may directly reduce survival through physical impairment, for example how improperly cured elastomer can migrate to organs where it can presumably impair organ function (Cabot et al 2021; McCarthy & Parris 2004). Even when no physical impairment occurs, marking may elicit behavioural responses immediately post-marking that have the potential to mediate downstream welfare and survival. Short-term behavioural responses to marking have been largely unexplored (but see Iannella et al 2017; Sapsford et al 2014), a knowledge gap which may overlook opportunities to improve welfare without abandoning methods, particularly for cases where the effects are transitory.

While most extant amphibians are anurans, caudates are disproportionately more threatened (57.3% caudates vs. 33.2%; IUCN 2023). Despite this, the majority of assessments of marking techniques have focussed on anurans (but see Ott & Scott 1999; Davis & Ovaska 2001; Kinkead et al 2006; McCarthy et al 2009). Here, we examined the impacts of two invasive marking techniques on the European alpine newt (*ichthyosaura alpestris*), a caudate species that is often the subject of numerous demographic and behavioural studies (recent examples: Bernabò et al 2023; Gvoždík 2022; Diego-Rasilla & Phillips 2021). The overriding aim of the study was to systematically evaluate the short-term effects of handling versus toe-clipping and VIE marking on newt behaviour. We achieved this aim through an experiment designed to discriminate between the effects of capture from the impacts of marking on newt activity, shelter use, and feeding.

## Methods

Alpine newts were collected from invasive populations in the UK and treated prophylactically with itraconazole (1 mg/L; Sporanox, Janssen-Cilag) to eliminate infections with *Batrachochytrium dendrobatidis* (Garner et al. 2009). Our previous work showed that post-treatment newt behaviour was still informative for comparative studies examining health and welfare (Daversa et al 2018). Newts (n=40, 32 female, 8 male) were weighed to the nearest 0.1 g, measured snout-to-vent (SVL) to the nearest mm and then individually housed in 5-litre plastic tubs (Really Useful Boxes, 340 × 200 × 125 mm), where they were given 5 days to acclimate. Each tub was divided in half, one half containing 1.5L aged tap water while the other was filled with autoclaved gravel (5-20 mm diameter). Cover objects (small PVC shelters) were embedded on gravel substrate and submerged in water. During acclimation and throughout the experiment, 1/3 of the tub water was replaced twice per week and debris removed from the aquatic side using sterile turkey basters to maintain sanitary aquatic conditions. We fed newts earthworms (*Lumbricrus terrestris*) after water changes (twice/week; total mass at each feed 0.4-0.5 g).

Experimental treatments were designed to discriminate between the effects of handling versus the effects of either marking protocol. Treatment #1 (placebo) involved capturing and holding newts in the hand for approximately 4 minutes before returning to the housing unit. Treatments #2 (toe-clipping) and #3 (VIE) were, respectively, hand-capture followed by complete removal of the middle digit of the rear right foot and return to housing unit, and hand-capture followed by injection of approximately 0.015 mL of elastomer in the ventral side of the thigh of the right rear leg and return to housing unit. Newts in the #4 treatment (control) were left undisturbed during the marking period. We balanced the sex ratio (8 females for every 2 males) across the four treatments, but within sex, newts were assigned randomly to treatments.

Newt behaviour was sampled for ten minutes immediately following return to the aquatic part of the housing unit. We recorded position (*aquatic* versus *terrestrial, hidden with head and greater than half the body under cover versus in view with head and greater than half the body exposed*) and activity (*active* versus *stationary*) for 30 seconds at 60 second intervals. We repeated focal sampling 48 hours and 7 days after the initial sampling. To eliminate treatment effects of time of day on sampling, we assigned one individual per treatment to one of ten sampling groups and collected behavioural data simultaneously across all 4 treatments in a sampling group. We recorded feeding behaviour immediately after handling/marking and once weekly after that for 3 weeks, as a yes/no event and as latency (time taken to show interest in food up to ten minutes after the food was added to the enclosure). We also recorded total consumption. The experiment was ended 21 days after marking.

For data analysis, we characterized three behavioural responses: the proportion of time active, proportion of time visible, and proportion of time aquatic. We ran Generalized Linear Mixed Models (GLMMs) in R (R Core Team, 2023; *lme4* package), one each for each response, to examine (a) how newt behaviour changed over the course of the experiment and (b) how handling and marking influenced newt behaviour. We used a binomial error structure for GLMMs and included a unique newt identification as a random effect to account for repeated sampling. We also included experimental treatment, observation number, and their interaction as fixed effects. We assessed the influence of the fixed effects on newt responses by performing likelihood ratio tests with a chi-square distribution, using the dropterm() function (*MASS* package). In cases where newt behaviour depended on the timing of observation, we also assessed the effect of experimental treatment on newt behaviour separately for each observation. To do so, we ran GLMMs with a binomial error structure and an observation-level random effect to account for overdispersion of the data (Harrison, 2014). These models included experimental treatment as a single, fixed effect. We compared model coefficients for the three treatment groups (placebo, toe clipping, VIE) against coefficients for the control groups to determine whether newts that were handled and/or marked behaved differently than controls.

To test for feeding latency, we first performed a ‘time-to-event’ analysis (i.e. survival analysis). The response variables used were (a) the time within the ten-minute observation period when newts were first observed feeding on worms (0 -10 minutes) and (b) an event status indicating whether newts fed within the 10-minute observation period (0 = no observed feeding, 1 = observed feeding).

For example, newts that were not observed feeding were assigned values of 10 for the observation period and 0 for the event status. Second, we ran generalized linear models (GLMs) to detect treatment effects on total consumption as a proportion of the total food provided, including experimental treatment as the fixed effect and again using a binomial error structure. We again used likelihood ratio tests with a chi-square distribution and the dropterm() function to examine the overall influence of experimental treatment on total worm consumption. We also compared model coefficients for the three treatment groups against coefficients for the control group to determine whether newts that were handled and/or marked consumed more or less than controls. We explored feeding trends over time by running GLMMs with a binomial error structure and including individual ID as a random effect to account for repeated sampling and interactive effects of treatment and sampling week. We tested for interactive and additive effects by performing likelihood ratio tests with a chi-square distribution and the dropterm() function.

## Results

The proportion of time that newts were active varied over observations (fixed effect of sampling event X^2^ = 37.28, p = < 0.001; Fig. 1a), though handling/marking did not generally influence the proportion of time that newts were active (treatment:sample interaction X^2^ = 3.89, p = 0.273; main effect of treatment X^2^ = 4.98, p = 0.174). However, examining activity patterns on a sample-by-sample basis revealed effects of all experimental treatments at a specific time point. Newts in all 3 handling/marking treatments were more active than newts in the control group immediately following handling/marking, an effect not detected in subsequent samples (Fig. 1a, Table 1a). Newt visibility varied across different observations (fixed effect of sampling event X^2^ = 43.36, p = < 0.001); yet again, handling/marking newts did not generally influence the proportion of time that newts were visible versus hidden in shelters (treatment:sample interaction X^2^ = 4.96, p = 0.175; main effect of treatment X^2^ = 4.26, p = 0.236). Examining visibility on a sample-by-sample basis did not reveal any event-specific effects (Fig 1b, Table 1b). Newt habitat use was influenced by handling/marking, depending on the sampling period and marking protocol (treatment:sample interaction X^2^ = 72.05, p = <0.001). Newts that were VIE-tagged spent proportionally less time in the water than control and other treatment newts at the last sampling (Fig 1c, Table 1c).

**Table 1.** Coefficients of models of newt activity, visibility, and habitat use on a sampling-by-sampling basis. The outputs shown are for generalized linear mixed models that included a kfixed effect of experimental treatment to determine whether newts that were handled/tagged behaved differently than newts in the control group. The models used a binomial error structure and included an observation-level random effect to account for overdispersion of the data. Behavioural differences between handled/tagged newts compared with controls is denoted by a p-value of <0.05 and are highlighted in bold.

**(a) Newt Activity**
| **Sample 1** | **estimate** | **SE** | **t** | **p** |
| --- | --- | --- | --- | --- |
| Placebo | 0.96 | 0.42 | 2.29 | **0.022** |
| Toe Clip | 0.87 | 0.42 | 2.06 | **0.039** |
| VIE | 0.88 | 0.42 | 2.09 | **0.037** |
| **Sample 2** |  |  |  |  |
| Placebo | -0.63 | 1.07 | -0.59 | 0.557 |
| Toe Clip | 1.15 | 1.00 | 1.15 | 0.251 |
| VIE | 0.72 | 1.00 | 0.72 | 0.471 |
| **Sample 3** |  |  |  |  |
| Placebo | 0.62 | 1.34 | 0.46 | 0.645 |
| Toe Clip | 0.86 | 1.34 | 0.64 | 0.521 |
| VIE | -0.38 | 1.41 | -0.27 | 0.786 |
| **(b) Newt Visibility** |  |  |  |  |
| **Sample 1** | **estimate** | **SE** | **t** | **p** |
| Placebo | 0.46 | 0.87 | 0.53 | 0.602 |
| Toe Clip | 1.01 | 1.01 | 1.00 | 0.326 |
| VIE | -0.40 | 0.74 | -0.54 | 0.591 |
| **Sample 2** |  |  |  |  |
| Placebo | 4.52 | 3.16 | 1.43 | 0.152 |
| Toe Clip | 0.24 | 3.64 | 0.07 | 0.948 |
| VIE | 2.70 | 3.48 | 0.78 | 0.438 |
| **Sample 3** |  |  |  |  |
| Placebo | 2.14 | 3.89 | 0.55 | 0.582 |
| Toe Clip | 1.54 | 3.55 | 0.44 | 0.664 |
| VIE | 1.44 | 3.49 | 0.41 | 0.679 |
| **(c) Newt habitat use** |  |  |  |  |
| **Sample 1** | **estimate** | **SE** | **t** | **p** |
| Placebo | -0.70 | 1.23 | -0.57 | 0.569 |
| Toe Clip | -1.84 | 1.09 | -1.69 | 0.091 |
| VIE | -1.12 | 1.16 | -0.96 | 0.337 |
| **Sample 2** |  |  |  |  |
| Placebo | 0.32 | 3.51 | 0.09 | 0.927 |
| Toe Clip | 0.17 | 3.54 | 0.05 | 0.961 |
| VIE | 1.01 | 4.04 | 0.25 | 0.802 |
| **Sample 3** |  |  |  |  |
| Placebo | 0.57 | 3.44 | 0.17 | 0.869 |
| Toe Clip | 0.34 | 3.32 | 0.10 | 0.918 |
| VIE | -19.05 | 4.45 | -4.28 | **<0.001** |

**Fig 1.**
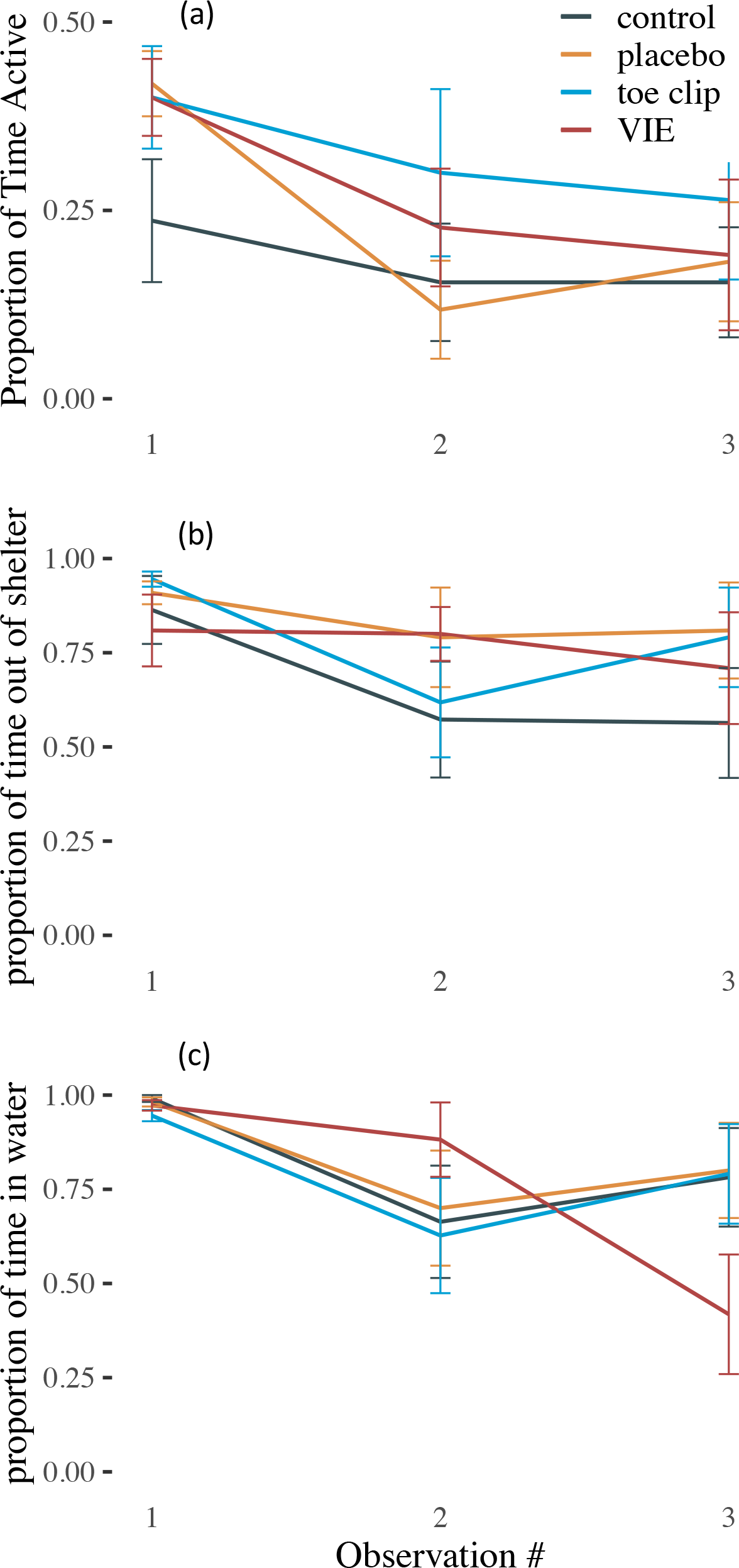
Temporal dynamics in newt behaviour following handling and marking procedures. Shown are the proportion of observations that newts were (a) active (i.e. moving), (b) outside of their shelter, and (c) in water as opposed to on land. Newts were sampled on 3 occasions: 1 -immediately after treatment, 2 - 48hr after treatment, 3 - 1 week after treatment.

Although there was no statistically significant treatment effect on feeding latency (including experimental treatment in Cox Proportional Hazards model only marginally improved fit, X^2^ = 6.60, p = 0.086), in qualitative terms, newts in the control group were, on average, quicker to start feeding after handling/marking than newts in treatment groups (Fig 2a). The above trends in feeding latency were not apparent after the first week (Fig 2b-d, Table 2b-d). There was a general treatment effect on the proportion of worms that newts consumed across the experiment (dropping treatment from the model reduced goodness of fit, X^2^ = 10.99, p = 0.012), but this effect was due to differences between newts receiving a placebo and newts that were toe-clipped (z = 3.20, p = .001; Fig 3). The proportion of worms that newts consumed in the three treatment groups generally did not differ to observed consumption in the control group (p > 0.05 in all cases, Fig. 3). Examining worm consumption on a weekly basis revealed interactive effects between marking/handing and the time of sampling (X^2^ = 10.61, p = 0.014), with effects largely driven by temporal changes in feeding by newts in the placebo group (Fig. 4). Newts in the placebo group (handled only) initially consumed fewer worms on average compared to subsequent observations (Fig. 4).

**Fig 2.**
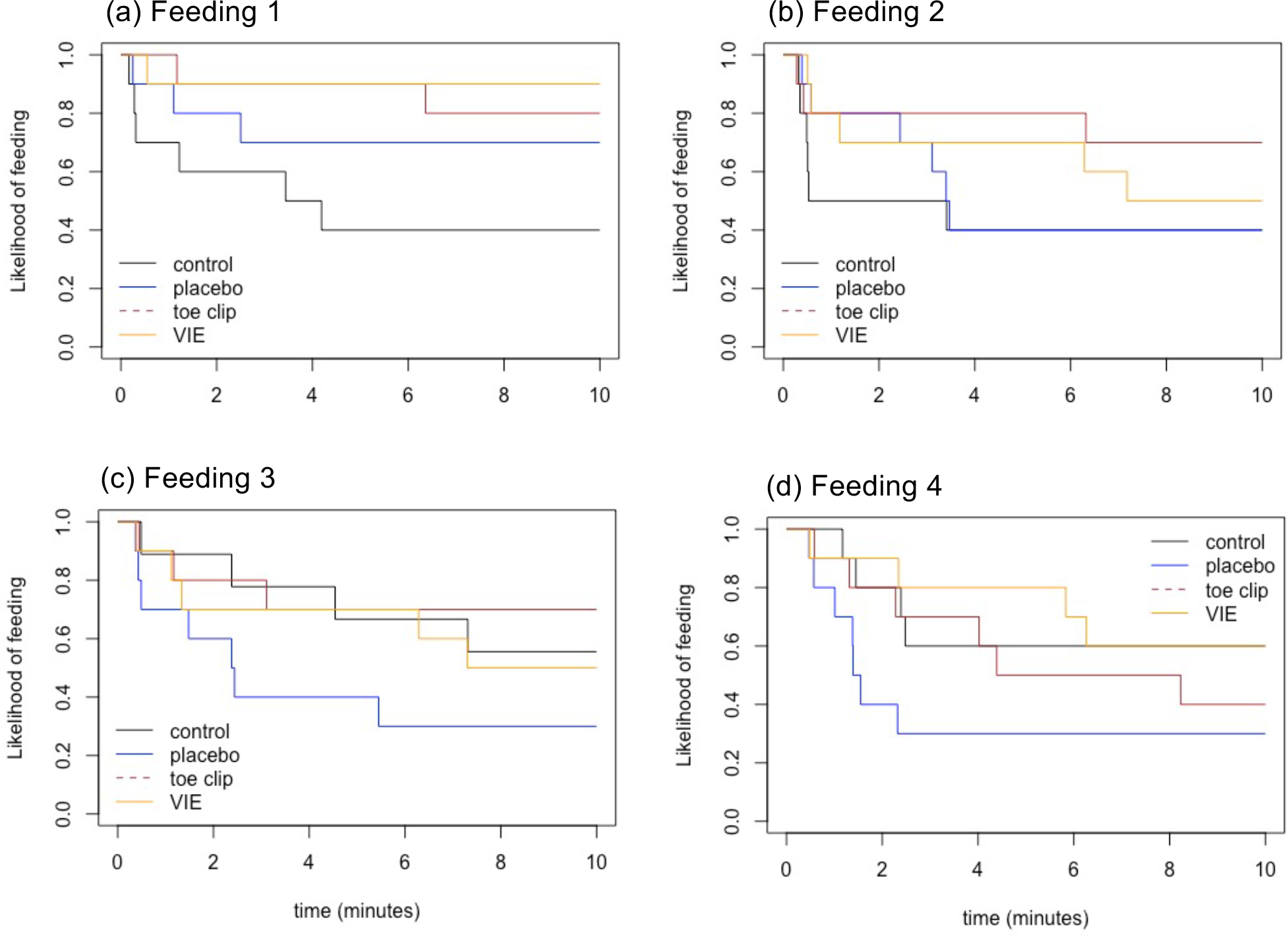
Time-to-event curves for feeding latency. Kaplan-Meir plots are showing the probability that newts were feeding at a given time during ten-minute observation periods performed weekly for four weeks. Observations began immediately after food was administered into newt tanks. Curves are shown for each of the four handling/tagging treatments.

**Fig 3.**
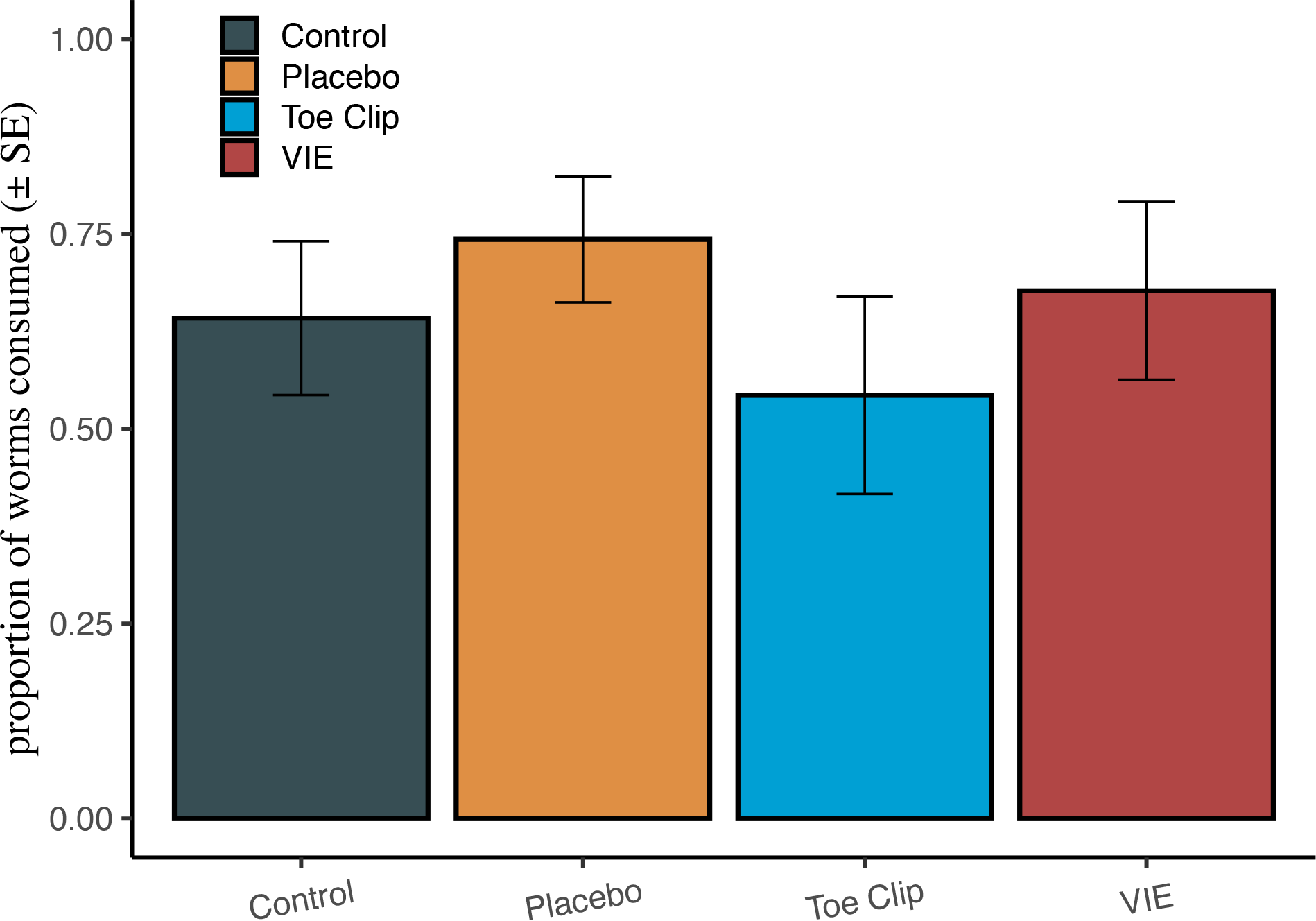
Food consumption is not influenced by handling or tagging. Shown is the mean proportion of worms consumed by newts that received different handling/tagging treatments. Newts were fed a total of 12 worms over the course of the experiment (4 feedings of 3 worms). Colors further denote the different experimental treatments and error bars show the standard error of the mean.

**Fig 4.**
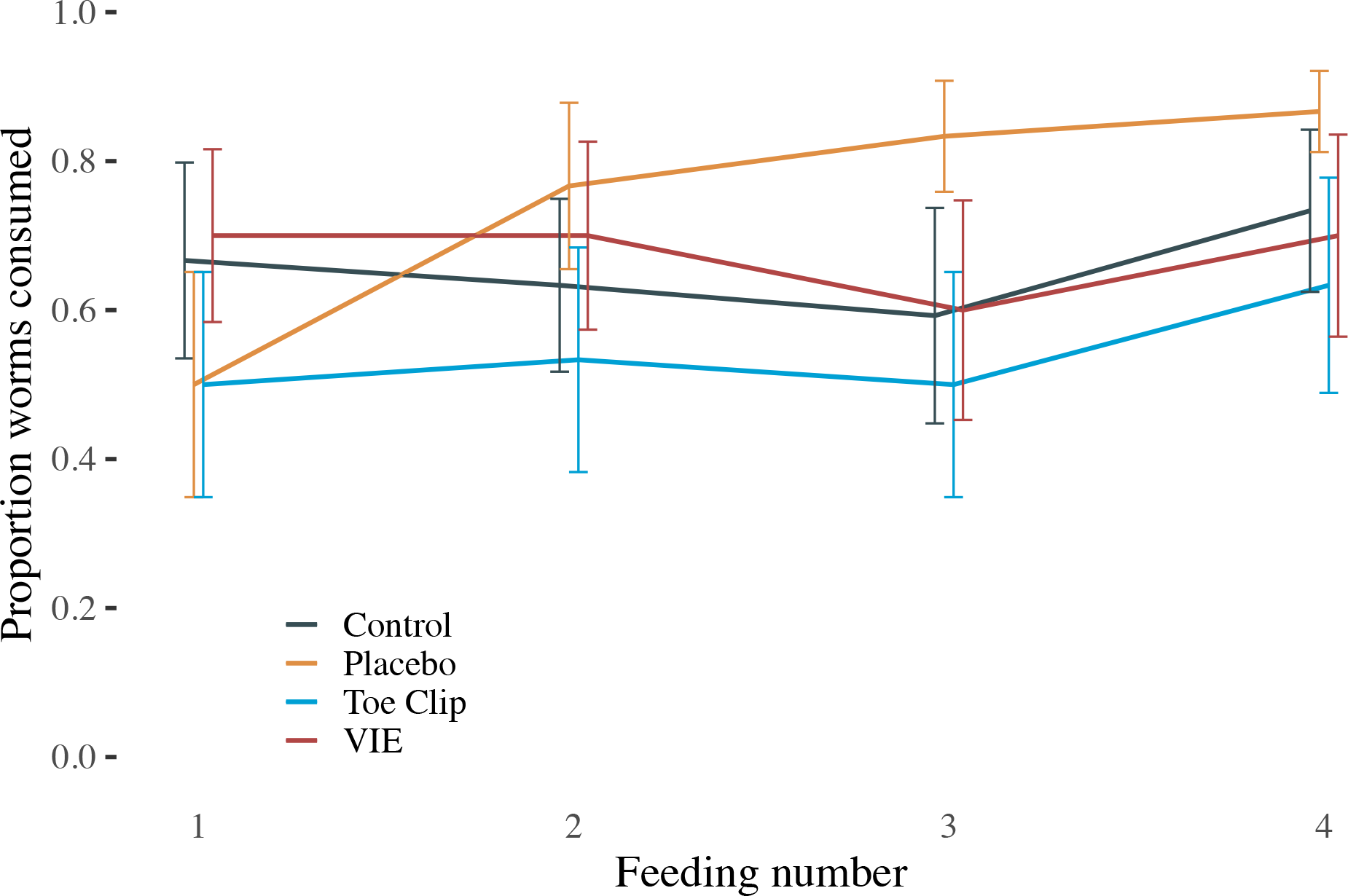
Newts receiving a placebo initially feed less. Shown is the mean proportion of worms consumed by newts on a weekly basis, with different lines distinguishing the specific marking/handling treatment received. Newts were fed three worms per feeding across four feedings (N = 12). The lines denote mean values within the treatment groups and error bars denote the standard error of the mean.

## Discussion

Assessments of the welfare impacts involved in the process of marking amphibians do not always discriminate between handling and marking (but see Oropeza-Sánchez et al 2020). This overlooks the directionality of the welfare assessment process, embedded in Soulsbury et al’s (2020) decision tree for marking wildlife. Here, the determination of both the necessity and welfare implications of capture precedes that of the impact of marking (Soulsbury et al 2020, Fig. 1). For a more meaningful assessment of the marking technique itself, then, the impacts of animal capture and restraint first need to be ascertained. Our study illustrates this, as we showed that the initial changes in activity by alpine newts elicited by handling and marking is largely attributable to the handling itself, and transient. Nevertheless, the initial increase in activity immediately after marking may not be in the best interests of the marked animal. Typically, reduced activity and immobility are amphibian responses to predator risk, and so increased activity possibly exposes animals to predators (Chapman et al 2017, Passos et al 2017). Yet, animals employ diverse antipredator behaviours depending on the context (Daversa et al 2021). In the context of handling by humans, the increased activity we observed is likely an antipredator escape behaviour initiated in response to what newts perceive as a predation attempt. This hypothesis is supported by studies of other wildlife that mount antipredator responses to human stimuli (Clinchy et al. 2016; Palmer et al. 2022). Interestingly, increased activity and feeding reductions were not associated with an increased propensity to seek refuge under cover objects or to move onto land.

The escape-like behaviours that we observed immediately after handling are characteristic of responses made in ‘fear’ and permit the hypothesis that human handling acts as a fear stimulus for newts (Daversa et al. 2021; Zanette & Clinchy, 2019). Fear-like responses in wild animals are well-studied and have widespread ecological consequences (Zannette and Clinchy 2020), and frameworks for understanding fear in wildlife hold value to welfare science. For example, fear-like escape behaviours involve physiological stress responses that cause temporary distress, heightened energetic demands, and internal damage (i.e ‘wear and tear’; Wingfield 2005, Sapolsky 2005). Stress responses involve a recovery period and may decrease investment in reproduction and feeding in caudates (Bliley & Woodley 2012; Moore & Miller 1984). In addition to relatively low initial feeding by newts in the placebo group, we observed increasing hesitancy to forage in 2 of the 3 handling treatments immediately after handling/marking, a trend that was strongest in the two treatments where marking was involved. Stress-associated inappetence may explain this hesitancy, an argument supported by evidence in badgers that human noise causes delayed feeding (Clinchy et al 2016). Alternatively, given that newts increased activity immediately after the marking/handling, they may have simply been too distracted to eat. In either case, the responses that we observed provide evidence that human handling induces fear-like responses in caudates, as they do in other animal groups (Clinchy et al. 2016; Palmer et al. 2022). However, our data also supports the conclusion that the effects on feeding are also transient, and any initial reduction in food intake can be compensated for.

There have long been calls for longitudinal studies in research of the ecology of fear (Daversa et al 2021). Longitudinal data is essential for measuring the duration of pain and/or distress and the how either might be behaviourally manifested. This study marks a step toward addressing these calls by tracking individual responses over time. Doing so, we found that behavioural responses to handling and marking were transient, providing hope that distress caused to animals is only temporary. The transient nature of the responses also suggests that newts may be able to develop tolerance to human handling that mediates fear and associated impacts to welfare. Nevertheless, repeatedly human handling can have covert physiological costs that accumulate over individual lifespans (Wingfield 2005) and should be explored more deeply to understand the extent of welfare impacts of handling and marking. Extending our experimental design to longer time periods, integrating measures of physiological rates (metabolic rate, hormone levels, etc.) and assessing physical damage from fear responses would mark a further step toward understanding how human handling and marking impact welfare in caudates and other wild animals.

## Acknowledgements

We thank the Biodiversity and Conservation MRes program at the University College of London and the Zoological Society of London (ZSL) for sponsoring this study. We also thank the students and staff in the Institute of Zoology, ZSL, for their kindness and moral support as we carried out the experiments.

## Ethics Statement

The experimental work and treatment with itraconazole were approved by the Zoological Society of London’s Ethics Committee before commencement and licensed by the Home Office (PPL 80/2466 to Garner).

## Author Contributions

EB, GR, and TG designed the experiment, with consultation from DD. EB and GR executed the experiment. DD performed the data analysis. TG wrote the initial draft of the manuscript, and all authors contributed to manuscript revisions.

